# Integument transcriptome profile of the European sea cucumber *Holothuria forskali (Holothuroidea, Echinodermata)*

**DOI:** 10.1101/2021.02.12.430961

**Authors:** Jérôme Delroisse, Marie Bonneel, Mélanie Demeuldre, Igor Eeckhaut, Patrick Flammang

## Abstract

In non-model organisms, Next Generation Sequencing (NGS) technology improve our ability to analyze gene expression and identify new genes or transcripts of interest. In this research, paired-end Illumina HiSeq sequencing has been used to describe a composite transcriptome based on two libraries generated from dorsal and ventral integuments of the European sea cucumber *Holothuria forskali (Holothuroidea, Echinodermata)*. A total of 43,044,977 million HQ reads were initially generated. After *de novo* assembly, a total of 111,194 unigenes were predicted. On all predicted unigenes, 32,569 show significant matches with genes/proteins present in the reference databases. Around 50% of annotated unigenes were significantly similar to sequences from the purple sea urchin *Strongylocentrotus purpuratus* genome. Annotation analyses were performed on predicted unigenes using public reference databases. These RNA-seq data provide an interesting resource for researchers with a broad interest in sea cucumber biology.

## 1. Introduction

Holothuroidea, also known as sea cucumbers, are worm-like soft-bodied marine organisms belonging to the echinoderm phylum and present worldwide. The class counts more than 1,600 species and several species are highly marketable as a food product in East Asian countries [1]. Nowadays, to continuously supply the high demand from the markets, new non-target species from the northern hemisphere are being fished and traded [2]. Sea cucumbers also possess a wide range of bioactive compounds that can potentially be used in the pharmaceutical industry [3, 4]. These compounds may present interesting biological activities (e.g. antioxidant, anticoagulant and wound healing, anti-inflammatory, antitumor or antimicrobial [5–7]). The sea cucumber *Holothuria forskali* is a common species found in the Eastern Atlantic Ocean and the Mediterranean Sea. This detritivore holothuroid of the family of Holothuriidae is found at shallow depth and is considered as a keystone species in its environment [8].

For non-model, or emerging model, marine organisms, Next Generation Sequencing technologies offer an opportunity for rapid access to sequence data and genetic information. Multiple echinoderm transcriptomes emerged in the literature in the last years [9, 10], bringing important molecular information on highly diverse biological processes such as development [11], regeneration [12], sensory perception [13], adhesion [14] or evolution [15]. In the present study, paired-end Illumina HiSeq sequencing technology has been used to generate an integument transcriptome of the sea cucumber *H. forskali*. More specifically, dorsal (i.e. bivium) and ventral (i.e. trivium) integuments were investigated separately as they were considered as functionally distinct. The transcriptome of the integument will be a valuable resource to better understand biological mechanisms occurring in this specific tissue and should positively impact future studies focusing on the species *H. forskali* but also on sea cucumbers in general. In particular, the distinction between ventral and dorsal integument libraries allows an emphasis on biological processes specific to both sides of the integument (e.g. the ventral integument includes tube feet involved in adhesion to the substratum, the dorsal integument is thought to be involved in sensory perception).

## 2. Data description

### 2.1. Animal collection and RNA isolation

Adult individuals of *H. forskali* were collected in the vicinity of the Marine Station of Banuyls-sur-Mer (France) in summer 2014. After dissection, the dorsal and ventral body wall were separated and the integument (i.e. dermis and epidermis) was cut into small pieces. The tissues were treated with fresh Trizol® solution and RNA extractions were performed according to the Trizol® manufacturer’s protocols. The quality of RNA extracts was checked using 1.2 M TAE agarose gel electrophoresis and spectrophotometric measurements using a Nanodrop spectrophotometer (LabTech International). RNA quality was finally assessed by size chromatography using an Agilent 2100 Bioanalyzer.

**Table 1.**
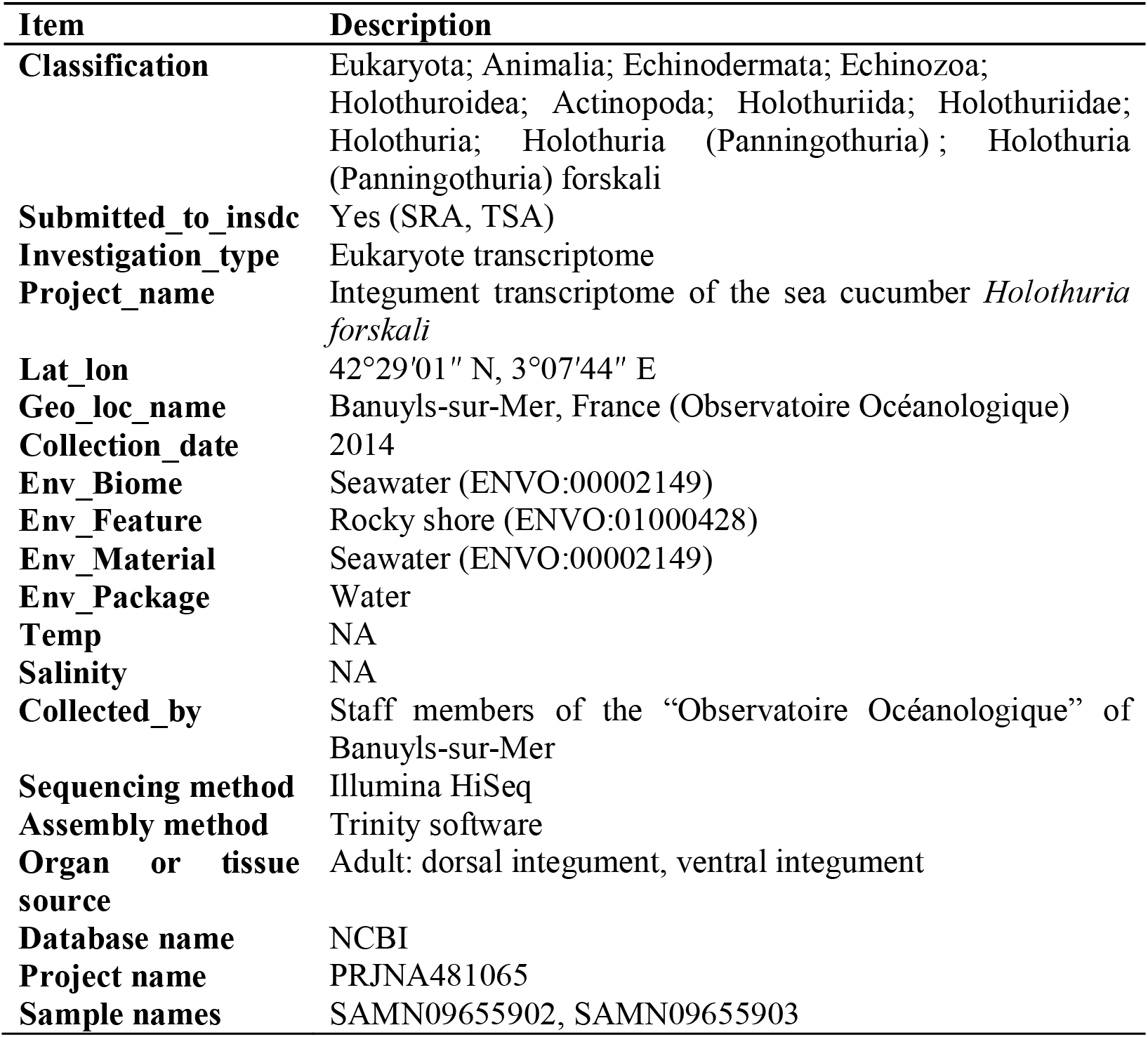
MIxS descriptors of the study

### 2.2. Library preparation and sequencing

Total RNA Samples were sent to a commercial sequencing service provider (Beijing Genomics Institute, Hong Kong). RNA samples were treated with DNase I and poly-(A) mRNAs were then enriched using oligo(dT) magnetic beads and fragmented into short pieces (around 200 bp). Random hexamer-primers were used to synthesize the first-strand cDNA using the short fragments as templates. DNA polymerase I was used to synthesize the second-strand cDNA. After purification, Double-stranded cDNAs were subjected to end reparation and 3’ single adenylation. Sequencing adaptors were ligated to the adenylated fragments, which were then enriched by PCR amplification. High-throughput sequencing was conducted using the Illumina HiSeq™ 2000 sequencing platform to generate 100 bp paired-end reads. Sequencing was performed according to the manufacturer’s instructions (Illumina, San Diego, CA).

### 2.3. Data processing and *de novo* assembly

The quality of the new transcriptomes was checked using the software FastQC (www.bioinformatics.babraham.ac.uk). An initial data cleaning was required to obtain clean reads that were used for the analyses. The cleaning step was performed by the sequencing service provider. It includes (i) the adaptor removal as well as (ii) the application of a filtering criterion to remove reads with more than 5% of unknown bases and low-quality reads (reads that comprise more than 20% low-quality bases, i.e. base quality ≤ 10). The Q20 percentages (i.e. base quality more than 20) were superior to 96,95% for both datasets.

After the initial cleaning step, the remaining 21,884,354 (ventral integument library) and 21,160,623 (dorsal integument library) clean reads were used to assemble the *H. forskali* integument transcriptome using the Trinity software (release 20130225) [16]. The following parameters were used: *seqType fq, min_contig_length 100, min_glue 3, group_pairs_distance 160, path_reinforcement_distance 95, min_kmer_cov 3.*

For the ventral transcriptome, 216,144 contigs were generated with an average of 377 base pairs (bp) while 187,773 contigs were generated for the dorsal transcriptome with an average length of 391 bp. The N50 (i.e. median contig size) was of 780 bp for the ventral contig set and 814 bp for the dorsal contig set.

Using paired end information and gap filling, contigs were further assembled into 111,194 unique sequences (i.e. non-redundant sequences or unigenes) with a mean length of 1,048 bp including 40,067 clusters and 71,127 singletons. Numerical data are summarized in Supplementary Table S1.

Unigenes were separated into clusters (similarity among overlapping sequences is superior to 94%) and singletons (unique unigenes). The clustering was performed using the TIGR Gene Indices Clustering (TGICL) tools (v2.1, parameters: *-l 40 -c 10 -v 20*) [17] followed by Phrap assembler (www.phrap.org, release 23.0, parameters : *repeat stringency 0.95, minmatch 35, minscore 35*). Various quality assembly criteria were evaluated such as (i) the contig/unigene size distribution and the (ii) of the read distribution when realigned to unigenes using SOAP aligner (Release 2.21, parameters: *-m 0 -x 500 -s 40 -l 35 -v 5 -r 1*) [18]. Length distribution of contigs and unigenes are presented in Supplementary Figure S1.A-D. In addition, more than 82% of transcriptome unigenes from the integument of *H. forskali* were realigned by more than 5 reads (Supplementary Figure S1.E).

### 2.4. Unigene annotation, classification and comparative gene expression

Transcriptome completeness was evaluated using BUSCO (v3.0.2) analyses on assembled unigenes (Supplementary Figure S2) [19]. Scores were calculated using Eukaryota_odb9 lineage data. BUSCO analyses showed that 91.1% complete BUSCO groups were detected in the final gene set (84.2% in the dorsal transcriptome, 77.3% in the ventral unigenes). In detail, out of the 303 evaluated BUSCOs from the Eukaryota dataset, only 8.3% were fragmented and 0.6% were missing.

Unigenes were compared to online databases NCBI non-redundant protein and nucleotide databases (NR/NT, www.ncbi.nlm.nih.gov, release 20130408), Swiss-Prot database (www.expasy.ch/sprot, release 2013_03), Kyoto Encyclopedia of Genes and Genome (KEGG, release 63.0) (www.genome.jp/kegg) and Cluster of Orthologous Groups (COG) of proteins (www.ncbi.nlm.nih.gov/COG, release-20090331) using BLASTx with a E-value threshold of 10^−5^, and in the nucleic-acid database (NCBI NT) by BLASTn with the same cutoff. When different databases returned inconsistent results, they were prioritized in the following order: NR, SwissProt, KEGG, COG. When a unigene did not align with any of the reference sequences, ESTScan was used to predict candidate coding regions and determine the direction of the coding sequence in the unigene [20].

On the 111,194 unigenes, 32,569 show significant matches with reference databases: 30,625 to NR, 10,397 to NT, 25,400 to Swiss-Prot, 22,813 to KEGG, 11,125 to COG and to 13,833 GO (Figure 1.A, Supplementary Table S2 for the complete unigene annotation table). The E-value distribution is presented in Figure 1.B. This E-value distribution of the “top matches” in the NR (NCBI) database showed that more than 57% of the mapped unigenes have strong homology (E-value < 1.0e^−30^), whereas 44% of the homologous sequences presented E-values ranging from 1.0e^−05^ to 1.0e^−30^ (Figure 1.B). The sequence similarity distribution indicates that 21% of the sequences have a similarity higher than 60% (Figure 1.C). Many unigenes were similar the genes found in the sea urchin *S. purpuratus* (Figure 1.D) as observed in other echinoderm-focused transcriptomic studies [11, 13].

**Figure 1.**
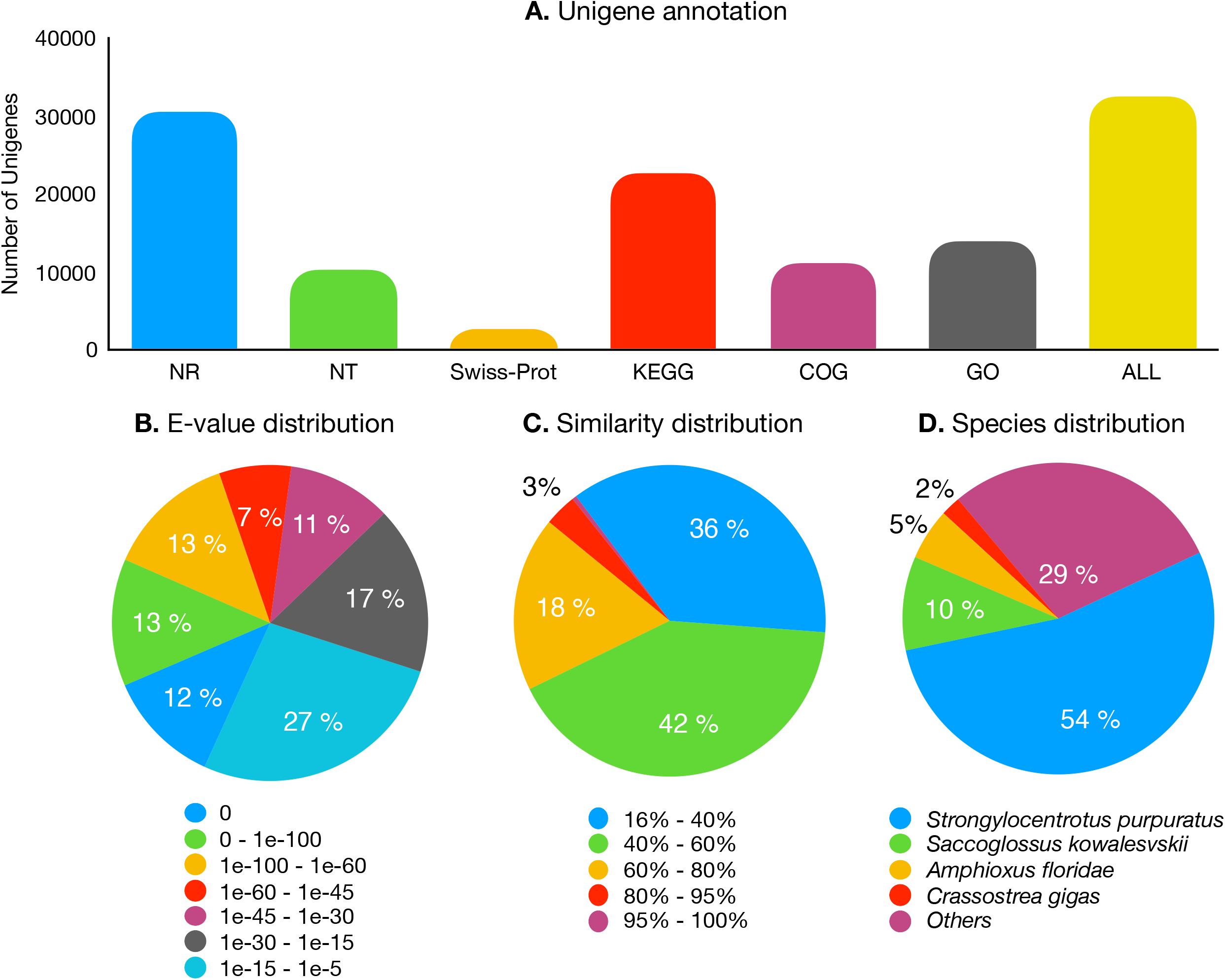
Annotation statistics of the integument transcriptome from *Holothuria forskali*. (A) Summary of the functional annotation of the unigenes for NR, NT, Swiss-Prot, KEGG, COG, GO databases. (B) “E-value distribution” of the top BLAST hits for unigenes (E-values < e^−5^). (C) “Similarity distribution” of BLAST hits of each unigene compared to the NR (NCBI) database. (D) “Species distribution” of the top BLAST hits for all unigenes.

To investigate the enzymatic diversity of both ventral and dorsal transcriptomes, additional annotation analyses were performed using the PRIAM database [21] implemented in the webtool FunctionAnnotator [22] using a stricter E-value threshold (cutoff of 10^−10^) (Supplementary Table S3).

On a total of 111,194 predicted unigenes, 30,104 were only found in the ventral integument transcriptome and 27,884 only in the dorsal integument transcriptome while 53,206 were detected in both transcriptomes. A comparative gene expression analysis was performed by mapping FPKM values (i.e. log_10_[FPKM value ventral integument transcriptome]) against log_10_[FPKM value dorsal integument transcriptome]), calculated for all predicted unigenes (Figure 2.A). However, it has to be stated that the transcriptomes have been generated in the purpose of new gene discovery and no biological or technical replication was performed as a part of the study. Based on a specific threshold (|log_2_[Fold change]| ≥1), 20,488 unigenes were found to be upregulated in the ventral integument transcriptome against 14,087 in the dorsal integument transcriptome (Figure 2.B).

**Figure 2.**
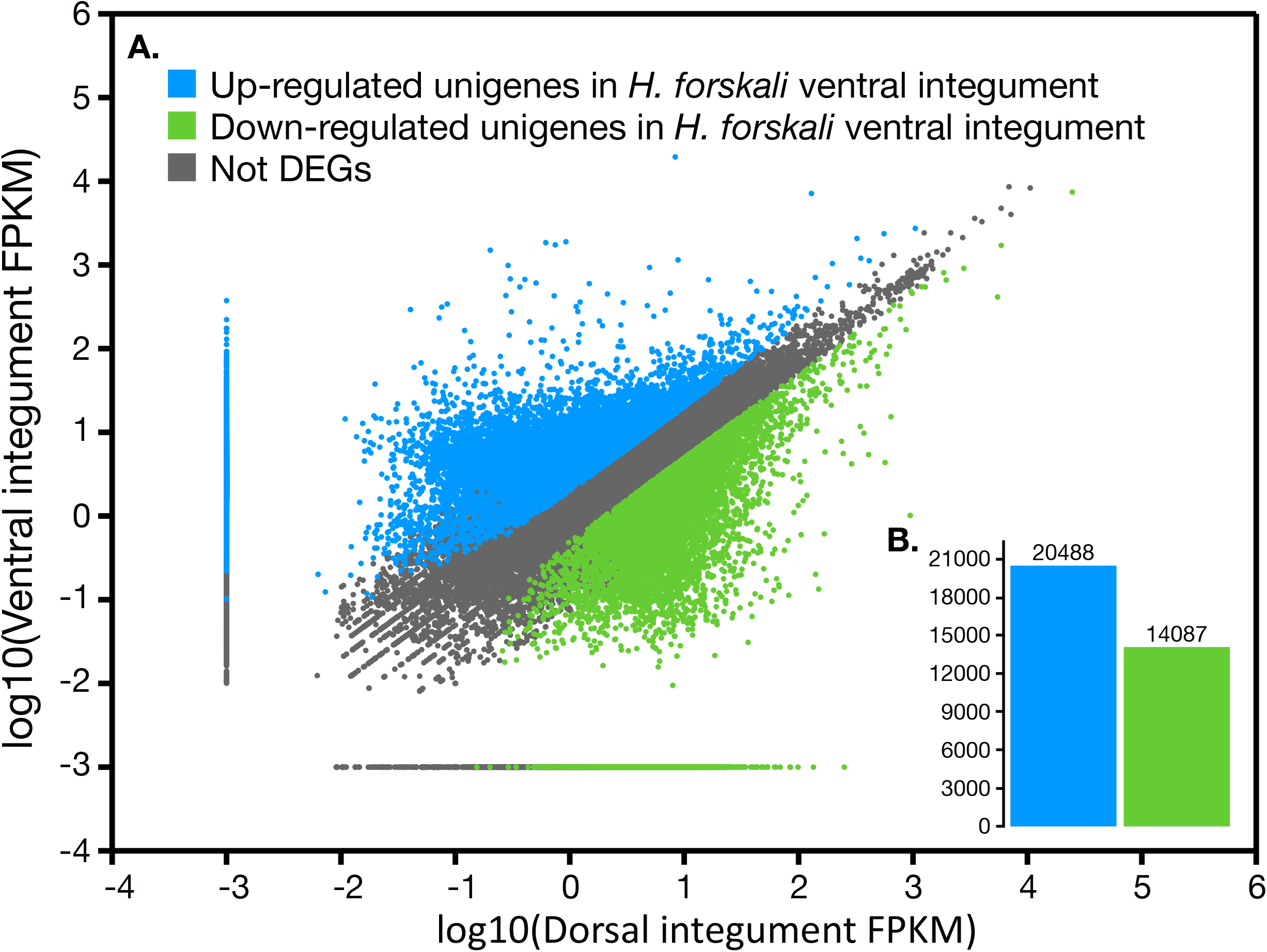
Comparative gene expression in H. forskali ventral and dorsal integuments. A. FPKM distribution of both transcriptomes. Upregulated and downregulated are color coded. Selection was based on the threshold: |log_2_[Ratio] |>= 1 (i.e. Log_2_[FPKM ventral + 0.1)/(FPKM dorsal + 0.1)] B. Comparison of the differentially expressed unigenes between the ventral and dorsal integument transcriptomes.

The 10 most expressed unigenes of both transcriptomes correspond to various subunits of the “cytochrome c oxidase”, a “thymosin beta-4”, an “alpha-1 collagen precursor”, a “ferritin” and several non-annotated unigenes (Supplementary Table S4). An “epidermal growth factor” is also specifically present within the 10 most expressed unigenes of the ventral transcriptome.

The 10 most differentially expressed unigenes (i.e. maximum or minimum log_2_[FPKM fold change]) in both transcriptomes are listed in the Supplementary Table S5. Most of the “10 most differentially expressed unigenes” with the highest expression in the dorsal transcriptome are not annotated. The 10 most differentially expressed unigenes with the highest expression in the ventral transcriptome correspond to a “rtoA-like”, an “hyalin-like”, a “farnesoic acid o-methyltransferase-like”, a “lactadherin-like”, a “rhamnose-binding lectin-like” and several non-annotated unigenes. Several of these actors are likely to be specifically expressed in tube feet and involved in tube foot adhesion such as the “hyalin-like” and the “farnesoic acid o-methyltransferases-like”. Hyalin proteins are fibrillar glycoproteins involved in cell adhesion and expressed, for example, in the sea-urchin *S. purpuratus* embryo [23]. Various “farnesoic acid o-methyltransferases-like” have recently been identified in footprint produced by sea star tube feet indicating a probable implication in the adhesive material elaboration [24].

For descriptive purposes, and based on PRIAM annotation, the 10 most differentially expressed predicted enzyme-coding transcripts are listed in the Supplementary Table S6. The most differentially expressed predicted enzyme-coding transcripts within the ventral integument transcriptome are several “ Protein-tyrosine-phosphatases”, a “Guanylate cyclase”, a “NAD+ ADP-ribosyltransferase”, a “Protein-serine/threonine kinase”, a “CDP-diacylglycerol--inositol 3-phosphatidyltransferase” and a “Creatine kinase”. Within the dorsal integument transcriptome, the most differentially expressed enzyme-coding transcripts are a “Lysozyme”, an “Exo-alpha-sialidase”, a “superoxide dismutase”, “DNA-directed RNA polymerase”, “Peptidylprolyl isomerase”, “Protein disulfide-isomerase”, “RNA helicase”, “Nicotinamide-nucleotide adenylyltransferase” and an “Amidase”.

## 2.5. Data accessibility

The Illumina derived short-read files are available at the NCBI Sequence Read Archive under the study accession number SAMN09655902 and SAMN09655903 (Bioproject N° PRJNA481065). This Transcriptome Shotgun Assembly project has been deposited at DDBJ/EMBL/GenBank under the accession GIPR00000000. The version described in this paper is the first version, GIPR01000000.

## Acknowledgments

J.D. and P.F. are respectively Postdoctoral Researcher and Research Director of the Fund for Scientific Research of Belgium (F.R.S.-FNRS). M.B. and M.D. are FRIA PhD students (F.R.S.-FNRS). The work was supported in part by (*i*) a PDR-WISD project (n° 29101409) from the F.R.S.-FNRS as well as the (*ii*) the FP7 European project BYEFOULING (Grant Agreement n° 612717). This study is a contribution from the “Centre Interuniversitaire de Biologie Marine” (CIBIM).

## Authors’ contributions

J.D. performed the experiments and data analyses. J.D., M.B., M.D., P.F. conceived and designed the experiments. J.D. wrote the first draft of the manuscript. All authors revised the manuscript.

## Supplementary data

**Supplementary Figure S1.** Distribution of contigs (A-B) and unigenes (C-D) in ventral and dorsal *Holothuria forskali* integument transcriptomes, respectively. The length of contigs and unigenes ranged from 200 bp to more than 3,000 bp. (E) Assessment of assembly quality using the distribution of unique mapped reads on the assembled unigenes.

**Supplementary Figure S2.** Transcriptome completeness evaluation on assembled unigenes using BUSCO.

**Supplementary Table S1.** Data description of the *Holothuria forskali* integument transcriptomes. A. Description of the sequencing output. Q20 percentage is the proportion of nucleotides with quality value larger than 20 in reads. GC percentage is the proportion of guanidine and cytosine nucleotides among total nucleotides. B. Summary statistics of transcriptome assembly.

**Supplementary Table S2.** Unigene annotation using NT, NR, GO, COG and KEGG databases (E-value threshold: 10^−5^).

**Supplementary Table S3.** Enzyme unigene annotation using PRIAM database (E-value threshold: 10^−10^).

**Supplementary Table S4.** The 10 most expressed unigenes in the ventral (A) and dorsal (B) integument transcriptomes of *Holothuria forskali* with their corresponding annotation.

**Supplementary Table S5.** The 10 most differentially expressed unigenes (i.e. maximum or minimum log_2_[FPKM fold change]) in the ventral (A) and dorsal (B) integument transcriptome of *Holothuria forskali* with their corresponding annotation.

**Supplementary Table S6.** The 10 most differentially expressed predicted enzyme-coding transcripts in the ventral (A) and dorsal (B) integument transcriptomes of *Holothuria forskali*.

## Compliance with ethical standards

### Ethical approval

All animal collection and utility protocols were approved by the Henan University of Science and Technology of Biology Animal Use Ethics Committee. Collections will be carried out in accordance with local and international laws. No special permits are needed for the marine invertebrate species used in this work and no ethics approvals are required for this study because research on echinoderms is not subject to ethics regulation. The animals used in our experiments were maintained and treated in compliance with the guidelines specified by the Belgian Ministry of Trade and Agriculture.

