## Supplementary figures and images for "Integument transcriptome profile of the European sea cucumber *Holothuria forskali (Holothuroidea, Echinodermata)*"

### Supplementary Figure S1

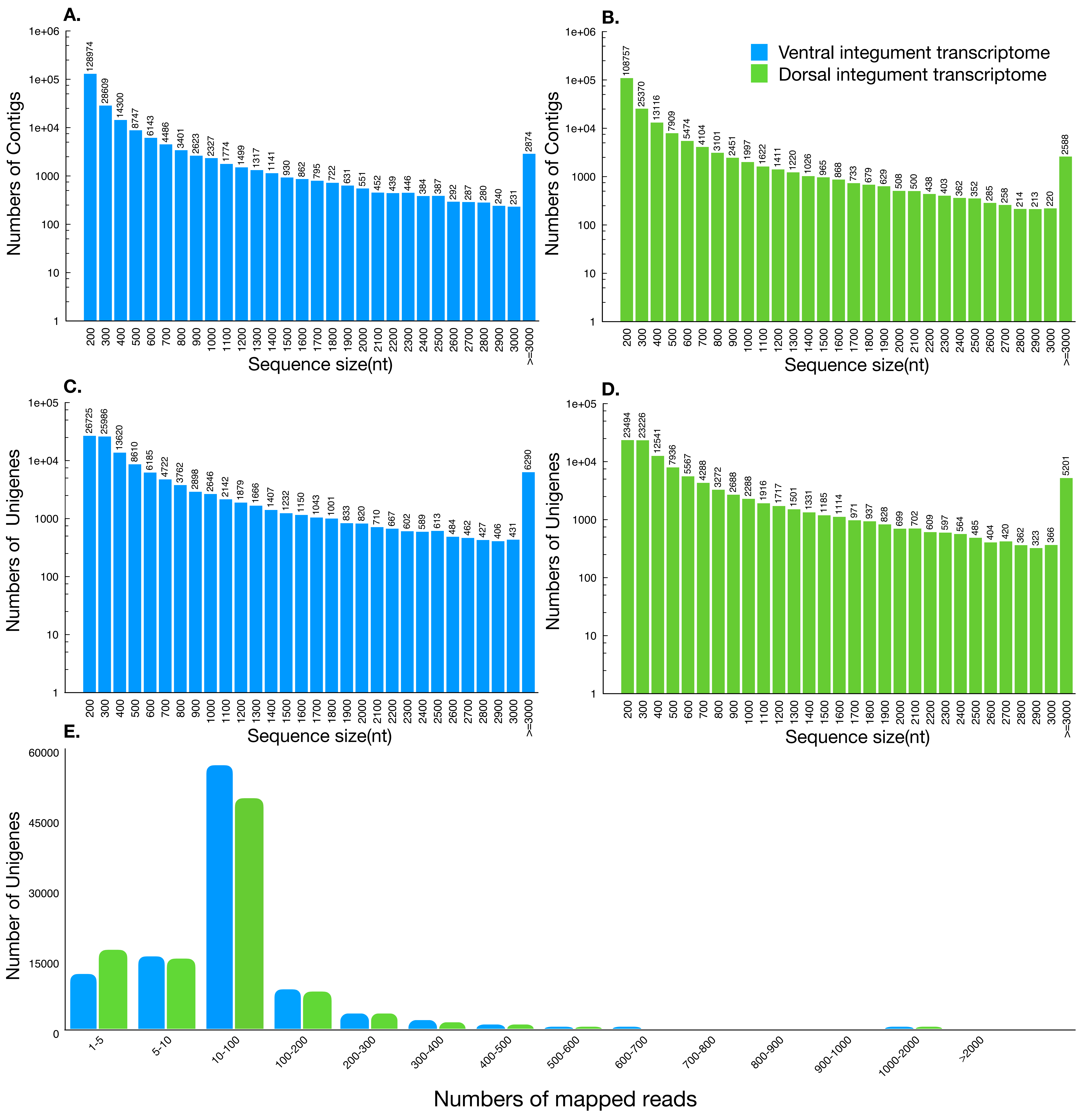

### Supplementary Figure S2

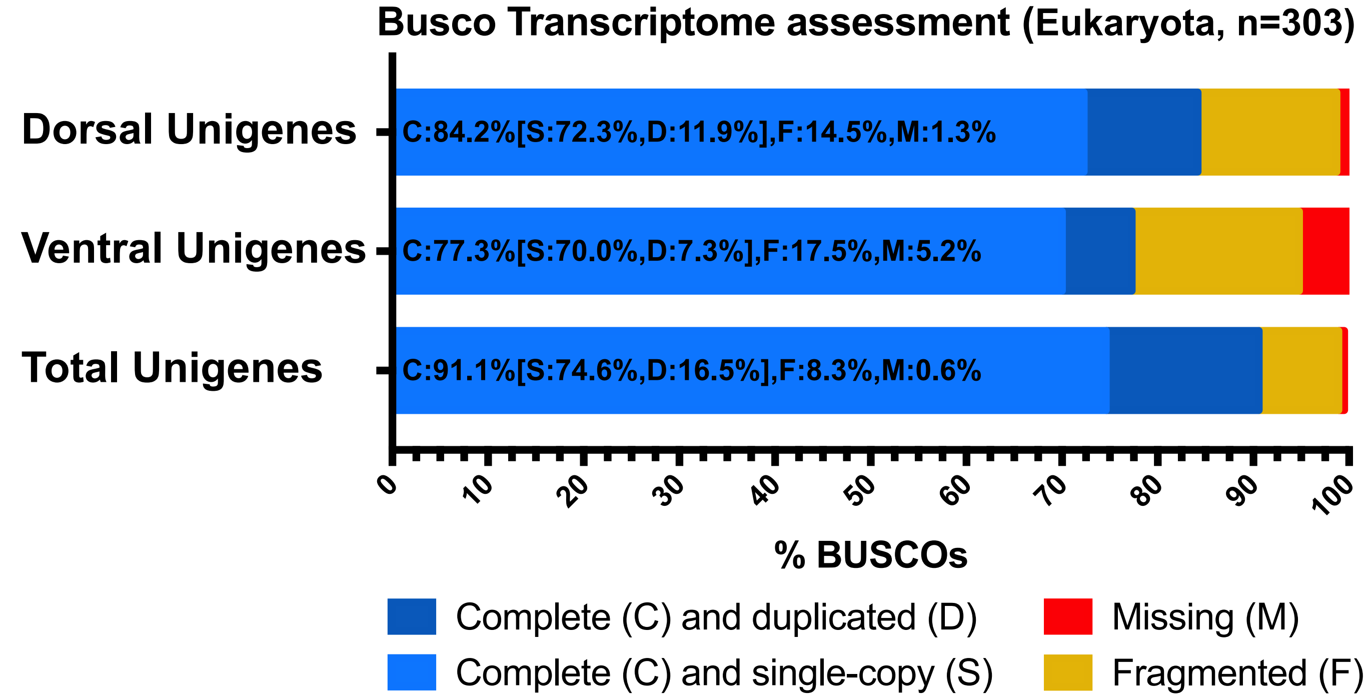
